# Structural Basis of the Transmembrane Domain Dimerization in the Activation Mechanism of TrkA by NGF

**DOI:** 10.1101/721233

**Authors:** María L. Franco, Kirill D. Nadezhdin, Sergey A. Goncharuk, Konstantin S Mineev, Alexander S. Arseniev, Marçal Vilar

**Affiliations:** Molecular Basis of Neurodegeneration Unit. Institute of Biomedicine of València (IBV-CSIC). C/ Jaume Roig 11, 46010 València, Spain; Shemyakin-Ovchinnikov Institute of Bioorganic Chemistry of the Russian Academy of Sciences, Moscow 117997, Russian Federation; Moscow Institute of Physics and Technology (State University), Institutskiy Pereulok 9, Dolgoprudny, Moscow Region 141700, Russian Federation

**Author notes:** Co-First author (María L. Franco, Kirill D. Nadezhdin).

## Abstract

Trk receptors are essential for the nervous system development. The molecular mechanism of TrkA activation by its ligand NGF is still unsolved. Recent data indicates that at endogenous levels most of TrkA is in an equilibrium monomer-dimer and the binding of NGF induces an increase of the dimer and oligomer forms of the receptor. An unsolved issue is the role of the transmembrane domain (TMD) in the dimerization of TrkA and the structural details of the TMD in the active dimer receptor. We found that TrkA-TMD can form dimers, identified the structural determinants of the dimer interface in the active receptor and validated this interface using site-directed mutagenesis together with functional and cell differentiation studies. Using *in vivo* crosslinking we identified a reordering of the extracellular juxtamembrane (JTM) region after ligand binding. Replacement of some residues in the JTM region with cysteine form ligand-independent active dimers and reveal a preferred dimer interface. In addition to that, insertion of leucine residues into the TMD helix induces a ligand-independent TrkA activation suggesting that a rotation of the TMD dimers could be behind TrkA activation by NGF. Altogether our data indicates that the transmembrane and juxtamembrane regions of the receptor play a key role in the dimerization and activation of TrkA by NGF.

## Introduction

Nerve growth factor (NGF) is a member of the mammalian neurotrophin (NT) protein family implicated in the maintenance and survival of the peripheral and central nervous systems (1–3). NGF is a dimer that interact with two distinct receptors, TrkA a cognate member of the Trk receptor tyrosine kinase family and the p75 neurotrophin receptor, which belongs to the tumor necrosis factor receptor (TNFR) superfamily of death receptors (4–6). TrkA signalling is essential for sensory and sympathetic neuron survival during development (7). Genetic mutations in the NTRK1 gene cause Congenital Insensitivity to Pain with Anhidrosis (CIPA)(8), and somatic mutations and chromosomal rearrangements generate aberrant protein fusions with constitutive kinase activation causing several types of cancer (9,10).

Despite all these important roles, the molecular mechanisms of TrkA activation have been poorly studied in comparison with those of other receptor tyrosine kinase (RTK) family members (11,12). The first three extracellular domains of TrkA consist of a leucine-rich region, LRR (Trk-d1) that is flanked by two cysteine-rich domains (Trk-d2 and Trk-d3). The fourth and fifth domains (Trk-d4 and Trk-d5) are immunoglobulin (Ig)-like domains, and these are followed by a 30-residue-long linker that connects the extracellular portion of the receptor to the single transmembrane domain and a juxtamembrane intracellular region that is connected to the kinase domain. TrkA is activated by nerve growth factor (NGF) a member of the neurotrophin family (3). The NGF binding domain is located in the Trk-d5(Ig2) domain (13) although other domains also participate in activation by NTs through an unknown mechanism (14,15).

Two models for TrkA activation are postulated; a ligand-induced dimerization of TrkA monomers and a ligand-induced conformational activation of pre-formed inactive dimers. The first model, which is based on the crystal structure of NGF with the ligand-binding domain of TrkA (13,16), assumed that the dimerization of TrkA is solely ligand-mediated and that receptor-receptor interactions are not present in the absence of its ligand. In the second model TrkA exists as a pre-formed inactive dimer suggesting receptor-receptor contacts in the absence of NGF (17,18). The most recent data using single-particle tracking (19) and FRET studies (20) suggests that TrkA is, at endogenous levels, predominantly monomer (80%) and NGF binding induces an increase and a stabilization of the TrkA dimers and the formation of oligomers, together with a conformational change leading to kinase activation. This mechanism of activation has been called the “transition model” (20) and postulates a dynamic transition from a monomer to an inactive dimer to a ligand-bound active dimer, suggesting that Trk receptors are activated through a combination of the two mentioned models.

Whatever the model, it is clear that dimerization of TrkA is required for its activation. Deletion constructs suggested that dimerization of TrkA in the absence of NGF is mediated by the transmembrane (TM) and by the intracellular domains (ICD) (20). In the case of the ICD this is supported by the crystal structure of the kinase domain of TrkA that showed the presence of dimers in the crystallographic unit (21,22). However the structural determinants of the TMD dimerization are not know and in this regard, it is important to understand the conformation of the TrkA-TMD dimer and identify the active dimer interface that may represent the functional state of the full-length receptor. In addition biochemical data supporting a conformational activation of TrkA pre-formed dimers are lacking.

In the present work we investigated the role of TrkA-TMDs in its activation, using several complementary structural and molecular biological approaches.

## Results

### Structural basis of TrkA transmembrane domain dimerization

It has been shown that the isolated TMDs of all human RTKs form dimers in bacterial membranes (23). In addition functional studies indicate that TMDs play an important role as a modulator of RTK homodimerization and kinase activation (reviewed in (23–25)). Switching between two dimerization modes of the transmembrane helix has recently been described as part of the activation mechanism of the EGF, VEGF and FGF receptors (26–29). However, to date the role of TrkA-TMD dimerization in TrkA receptor activation has not been studied in detail. Here we used two different approaches to characterize the dimerization of the TrkA-TMD. First, to study TrkA-TMD dimerization in a biological membrane context we used ToxRED, a genetic bacterial system, to assess TrkA transmembrane dimerization using ToxCAT as a reporter system (30,31). The TrkA-TMD self-associated in biological membranes in this system (Figure 1A). To determine the TrkA amino acid residues implicated in this dimerization we assayed the effect in this assay of individual mutation of each of the small residues (Ala, Gly or Ser) in the TMD sequence to a bulky residue such as isoleucine. This approach to disruption of dimerization was similar to that used in classical studies of glycophorin-A, in which the mutation G83I impaired its TMD dimerization (31). In our study, less TMD dimerization was observed in the S419I and G423I mutants compared to wt (Figure 1B), implicating the sequence S_419_xxxG_423_ as part of the TrkA-TMD dimerization motif in bacterial membranes.

**Figure 1.**
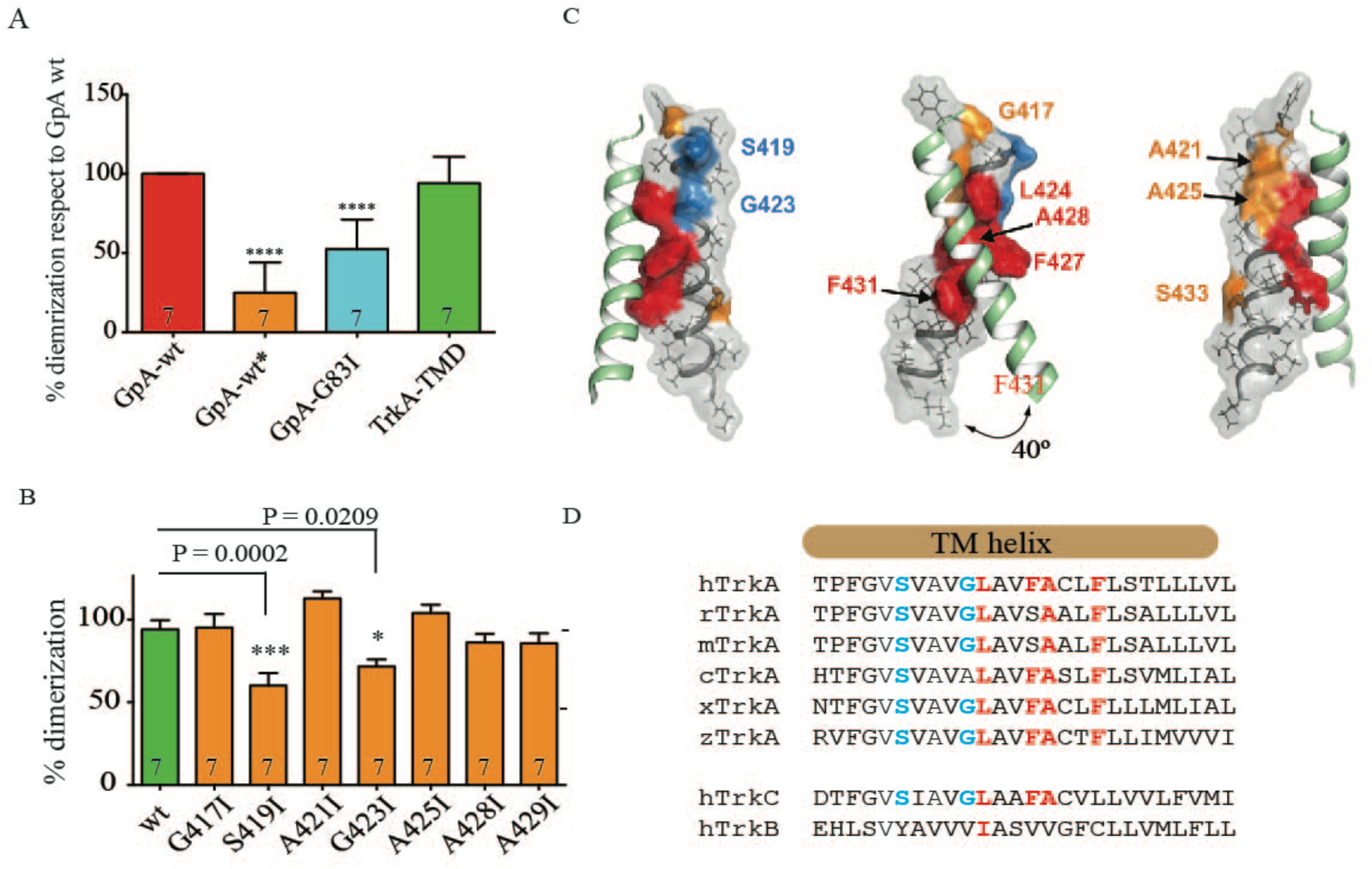
Analysis of the structure of TrkA transmembrane domain dimers in the ToxRED system and in DPC micelles. A) ToxRed system analysis and quantification of the percentage of dimerization of the rat TrkA-TMD normalized to glycophorin A (GpA) wt (100%). GpA-wt* refers to an inactive ToxR that cannot induce the RedFP. B) Percentage of dimerization of rat TrkA-wt and TrkA-TMD mutants normalized to dimerization of GpA wt (100%) in the ToxRed system. (In A and B results represent the average of at least 10 colonies analyzed per experiment. Plotted is the result of at least seven independent experiments). Error bars represent the standard error calculated from the seven independent measurements. Statistics were performed using two-way ANOVA analysis and Dunnet’s multiple comparison test using GraphPad software. P values of conditions significantly different from wt are shown on top of the error bars (**** P<0.0001). C) Schematic representation of the NMR spatial structure of the human TrkA-TMD dimer in lipid micelles, from different angles. The surface of one TMD helix is represented with the S_419_xxxG_423_ motif in blue and the NMR dimer interface in red. The TMD of the other monomer within the dimer is represented as ribbons, and side chains are indicated as sticks. The location of the NMR dimer interface LxxFAxxF on the NMR structure is colored red and other mutated residues are colored orange. The PDB accession code is 2n90. D) Alignment of the amino acid sequences of the transmembrane helical domain of TrkA from different species (h, homo sapiens, r, ratus norvegicus, m, mus musculus; c Gallus gallus; z Danio rerio; x Xenopus laevis). The residues participating in the NMR dimer interface are highlighted in red, and residues participating in dimerization based on the ToxRED analysis are highlighted in blue.

To obtain a structural insight into TrkA-TMD dimerization we solved the structure of human TrkA-TMD dimers in lipid micelles using NMR. For this study, the human TrkA-TMD was produced in a cell-free system (see Experimental Procedures) as previously described (32). When the peptide is solubilized in dodecylphosphocholine (DPC) micelles at a lipid-to-protein molar ratio (LPR) of 50:1 the TrkA-TMD is in equilibrium between monomeric, dimeric and other oligomeric states. The ratio of these states varies as the LPR value is altered (Figure S1A). We then titrated TrkA-TMD in DPC micelles using a standard technique in our laboratory (33) to measured the standard free energy of dimerization (ΔG_0_) (Figure S1B). The ΔG_0_ value obtained (−1.9±0.2 kcal/mol) suggested that the TrkA-TMD dimer is quite stable compared to the TMD dimers of other RTKs. Thus, although its dimerization energy is weaker than that of the VEGFR2 dimer (ΔG_0_ = − 2.5 kcal/mol in DPC) (34), it is stronger than that of FGFR3 (ΔG_0_ = −1.4 kcal/mol in DPC/SDS 9:1 mixture) and ErbB4 (ΔG_0_ = −1.4 kcal/mol in DMPC/DHPC 1:4 <u>bicelles</u>) (35). The ^15^N-HSQC spectrum of ^15^N labeled-TrkA-TMD (Figure S2) contained the expected number of cross-peaks, and the good quality of the spectra allowed solving of the structure of the dimer in DPC micelles (Figures 1C, Figure S3, Figure S4, Figure S5 and Table S1). The α-helical region of the TrkA-TM dimer starts at G417, ends at N440, and is ~38 Å in length (Figure S3). The crossing angle of the TrkA-TM helices is 40°, and the minimal distance between two monomers is 8.8 Å (Figure 1C and Table S1). The hydrophobicity plot and contact surface area of the dimer is shown in Figure S5. The dimerization interface lies along the sequence motif L_424_xxF_427_A_428_xxF_431_ (Figure 1C and Figure S5) that is conserved in the TrkA-TMD of several species and also in TrkC but not in TrkB (Figure 1D).

These analyses of TrkA-TMD dimerization suggested the existence of two possible dimerization motifs; S_419_xxxG_423_ and L_424_xxF_427_A_428_xxF_431_, which are located on opposite faces of the TMD helix (Figure 1C). The latter motif was found in analysis of the dimerization of TMDs in micelles using a direct approach and in the context of the isolated domain, while the former motif was observed in analysis of the dimerization of TMDs in real membranes, using an indirect approach and in the context of a chimeric protein. Thus, the biological relevance of each motif can be questioned, as the presence of large extracellular and intracellular globular domains or the lipid environment of the plasma membrane may favor or hinder a specific interaction interface (36). We therefore used different functional assays to verify the found dimer interfaces in the context of the full-length receptor.

### Functional identification of the dimer interface upon NGF stimulation

The state of full-length TrkA was followed by assay of three different aspects of its activity: dimerization of the receptor, phosphorylation of intracellular tyrosine residues and neurite differentiation of PC12nnr5 cells. To investigate the dimerization of TrkA, we individually mutated most of the N-terminal residues of the rat TrkA-TMD to cysteine (Figure 2A), expressed these constructs in Hela cells, and then measured the amount of cross-linked species. To facilitate cross-linking via these cysteine residues in the transmembrane domains we used oxidation with molecular iodine (I_2_) as previously described (37). Such oxidation allows the formation of a disulfide bond between two close cysteine residues inside the lipid bilayer. Plasma membrane fractions from cells expressing different single-cysteine mutants were incubated in the absence or presence of NGF, together with molecular I_2_ and were then analyzed by non-reducing SDS-PAGE and western immunoblotting. As shown in Figures 2B and 2C the mutants F416C, G417C and A421C formed covalent dimers in the absence of NGF. Upon NGF stimulation the amount of F416C, G417C and A421 dimers increased and a dimer was newly seen in the V418C mutant (Figures 2B and 2C). The residues G417, F416 and A421 are very close to each other in the NMR structure (Figures 2D and 2E), supporting the NMR-derived conformation as the active state of the TrkA dimer.

**Figure 2.**
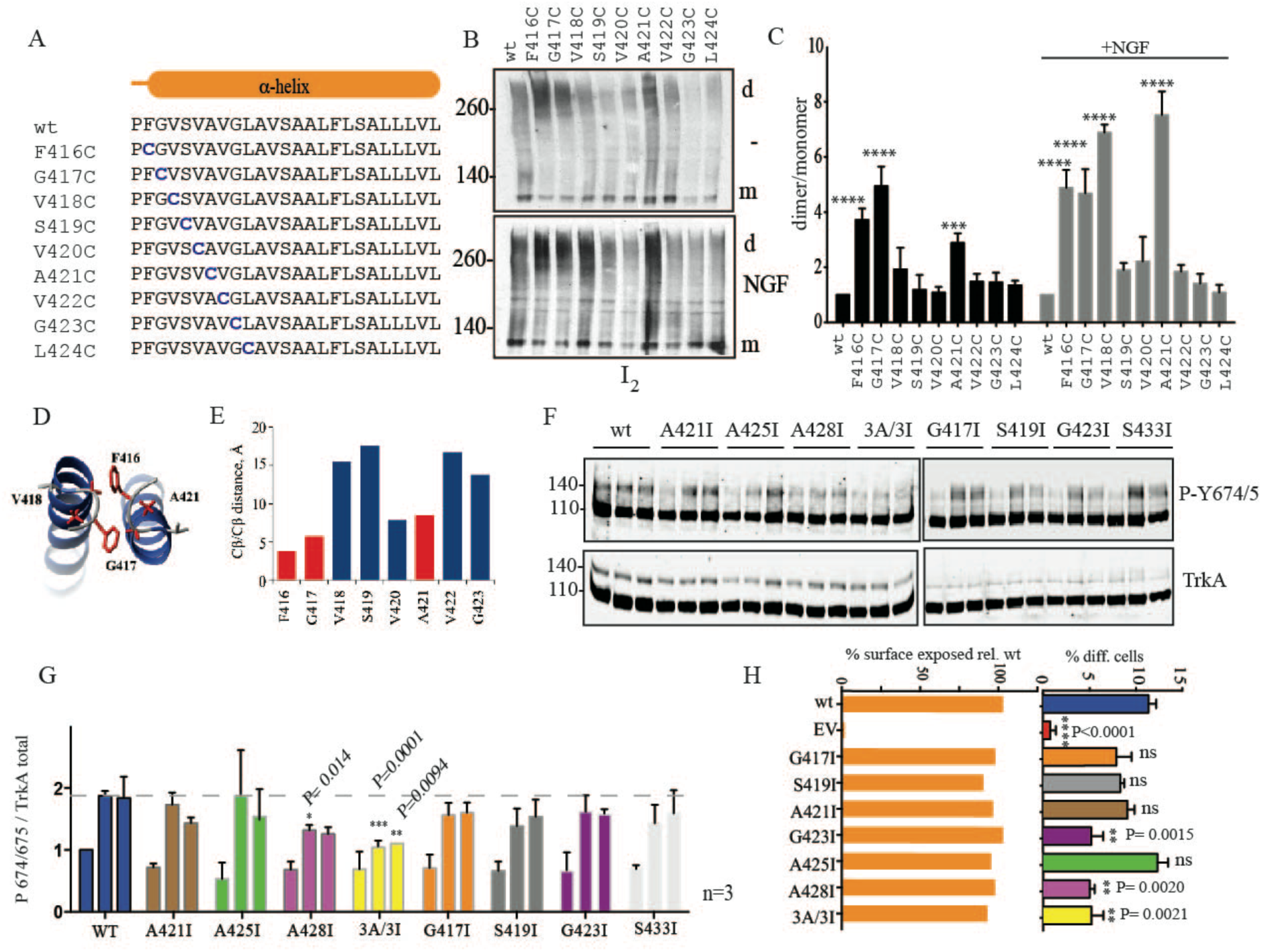
Functional identification of the two dimer interfaces in the TrkA-TMD. A) Amino acid sequence of the transmembrane domain of rat TrkA showing the location of the single cysteine residue substitutions. B) Western blot of Hela cells transfected with the indicated TrkA-TMD constructs and analyzed using non-reducing SDS PAGE, showing the formation of covalent cysteine dimers (d) arising from monomers (m) crosslinked using molecular iodine (I_2_) in the presence or absence of NGF. C) Quantification of the data in B) derived from at least three independent experiments. Bars represent standard error of the mean. Statistics were performed using two-way ANOVA analysis and Dunnet’s multiple comparison test using GraphPad software. (**** P<0.0001, *** P=0.0004). D) Structure of the TrkA-TMD dimer showing the location of the indicated residues in red. E) Cβ-Cβ distance between the indicated residues in the TrkA-TMD. F) Hela cells transfected with the indicated TrkA constructs were stimulated with NGF (10 ng/mL) for 0, 5 or 15 minutes. The immunoblots show the activation (autophosphorylation) of TrkA as detected using an antibody specific for TrkA P-Tyr674/675. The levels of total TrkA are shown below each blot. Molecular weight markers are shown at left. G) Quantification of the data in F) derived from at least three independent experiments. Bars represent standard error of the mean. Statistics were performed using two-way ANOVA analysis and Dunnet’s multiple comparison test using GraphPad software. P values of conditions significantly different from wild type are shown on top of the error bars. H) Cytometric analysis of the expression of the TrkA mutants at the plasma membrane (orange bars) and quantification of the percentage of PC12nnr5 differentiated cells (colored bars) transfected with the indicated TrkA constructs together with GFP. EV, empty vector. The percentage of the total GFP-positive transfected cells with a neurite twice as long as the length of the cell body was quantified. Bars represent the standard error of at least three independent experiments. Statistical analysis was performed using ordinary one-way ANOVA, using Bonferroni’s multicomparison test compared to wt. The P values of significant differences are shown. ns, not significant.

To further study the significance of the found interfaces, we mutated the small residues Ala, Gly and Ser within this region to the bulky Ile residue and we then assayed TrkA activation upon NGF stimulation (Figures 2F and 2G). The rationale behind this approach was that the mutation of a small residue to a bulky one on the relevant interface would prevent the formation of the active dimeric state by inducing steric clashes, and would, therefore, reduce TrkA activation. To perform this assay, we transfected Hela cells that do not express endogenous TrkA with these mutants, stimulated these cells with non-saturating concentrations (10 ng/mL) of NGF, and then assayed TrkA activation by analysis of TrkA autophosphorylation using western blotting. Upon transfection, two TrkA electrophoretic bands are present in the TrkA immunoblots of Hela cells; a lower band (approx.110 kDa) of intracellular immature TrkA that has not completed Golgi-mediated processing of high-mannose N-glycans (38) and an upper band (approx.140 kDa) with mature sugars that is expressed in the plasma membrane. Exposure to NGF substantially increased the phosphorylation of the upper TrkA band as assessed by blotting with a phospho-specific antibody against the phospho-tyrosine residues of the activation loop, Tyr674 and Tyr675 (P-Y674/5). This autophosphorylation was quantified to follow TrkA activation. Constitutive (t=0, no NGF added) and ligand-dependent phosphorylation of plasma membrane-localized TrkA after 5 and 15 minutes were measured. Since overexpression of TrkA induces ligand-independent autophosphorylation, we first transfected the Hela cells with increasing concentrations of TrkA to determine a TrkA level that could still be detected but that displayed no autophosphorylation in the upper band in the absence of NGF (Figure S6). It is noteworthy that all mutants are expressed at the plasma membrane as evidenced by both immunofluorescence localization in the absence of Triton X-100 using an antibody against an epitope in the TrkA N-terminus (Figure S7) and by flow cytometry (Figure 2H). Out of the seven single-point mutants tested only the A428I substitution demonstrated a pronounced inhibitory effect on receptor autophosphorylation. A428 is the only small-chain residue that is found deep in the dimerization interface of the TrkA TMD conformation determined using NMR, which further supports the relevance of the obtained NMR structure. The inhibitory effect of A428 substitution on receptor activity was further enhanced when all three Ala residues that are at least somehow involved in the TMD dimerization in the NMR-based structure, A421, A425 and A428, were simultaneously substituted (TrkA-3A/3I).

Lastly, we studied the effect of the same mutations on the NGF-induced differentiation of transfected PC12nnr5 cells (Figure 2H). Again, the A428I mutant displayed substantial inhibition of this TrkA activity. Unexpectedly, although the mutation S419I had no effect on TrkA activation by NGF in these two assays, the mutation G423I did have an impact on cell differentiation (Figure 2H).

The combined results of the functional assays agree regarding the importance of the NMR-derived TMD structure for TrkA activation. The critical role of the residue A428 in TrkA activation by NGF together with its location deep in the dimer interface indicates that the NMR-based structure corresponds to the mode of TMD interaction in the NGF-induced active state of the receptor

### Generation of constitutively active mutants of TrkA

Stimulation of Hela cells transfected with TrkA-wt with NGF induces the formation of TrkA dimers that are crosslinked with BS3 (Figure 3). While it is known that TrkA dimers are formed in the absence of NGF, no cross-linking was observed without the ligand, which suggests that ligand binding is accompanied by changes in the conformation of the extracellular part of the protein. As BS3 reacts only with free amines (the side-chains of Lys residues or a free N-terminus) we searched for possible sites in TrkA that might have caused the observed crosslinking. According to the crystal structure of the TrkA/NGF complex (15) (Figure 3A), there are no lysine residues in the TrkA-ECD that are located in a position where crosslinking of the side chains of Lys residues of two monomers could occur. Since BS3 does not cross the plasma membrane, and since we used the full-length TrkA receptor in our assays, we wondered if the observed BS3 cross-linking was mediated via crosslinking of K410 and K411 in the extracellular juxtamembrane region (eJTM) of TrkA (Figure 3A) since this region is not observed in the crystal structure (16). To verify this hypothesis, we mutated both K410 and K411 to Arg and repeated the initial experiment using HEK293 cells transfected with this TrkA-KK/RR construct (Figure 3B). No BS3-induced TrkA crosslinking was observed in the TrkA-KK/RR-transfected cells suggesting that NGF binding brings this region of the eJTM into close proximity.

**Figure 3.**
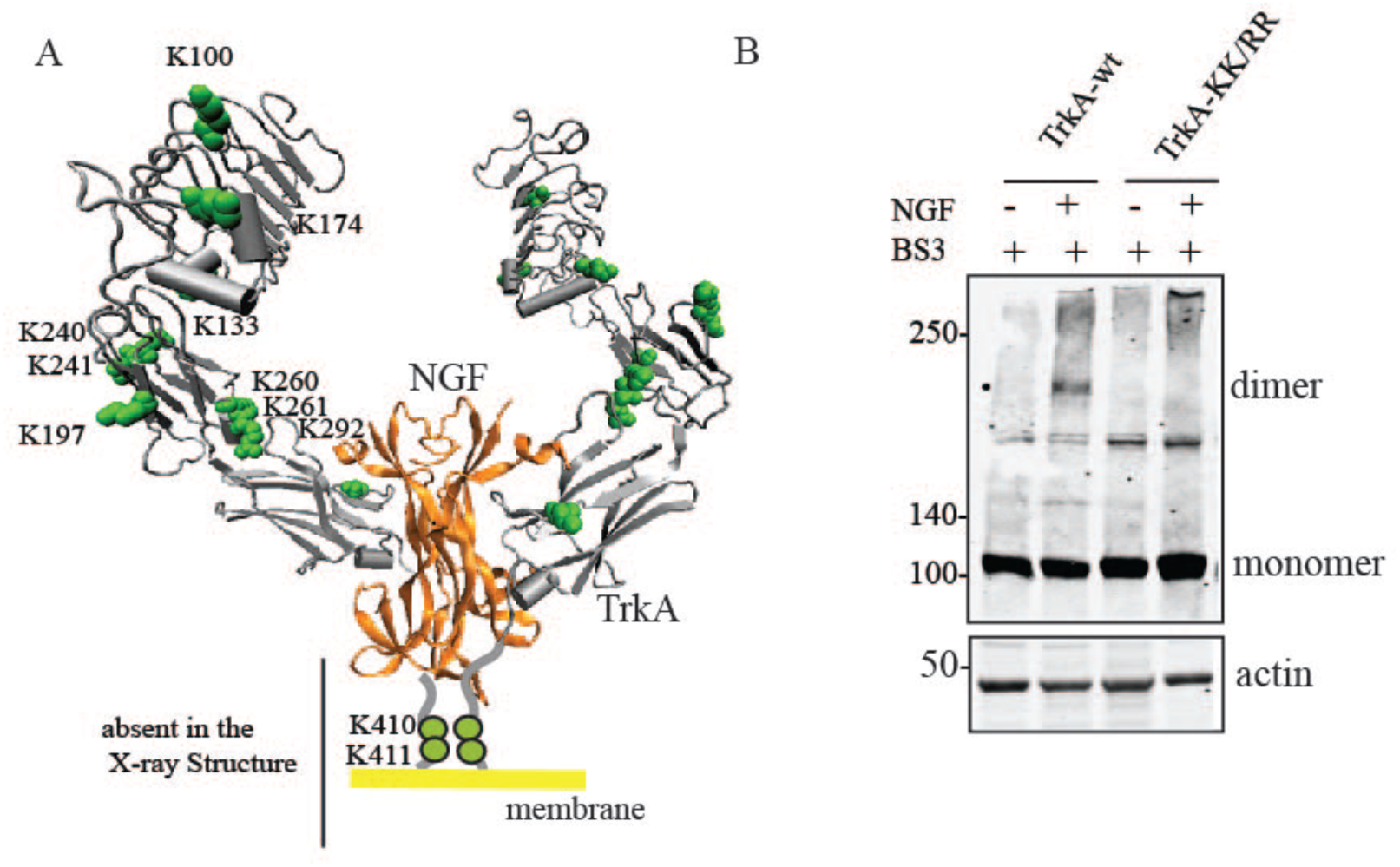
K410 and K411 of eJTM are crosslinked with BS3 upon NGF binding. A) Location of the Lys residues in the crystal structure of the rat <u>TrkA</u>-ECD/NGF complex (PDB: 2IFG) (Wehrman et al., 2007). The Lys residues are shown as green circles are located in the extracellular juxtamembrane region of the TrkA-ECD (in gray). B) Western immunoblots of lysates of Hek293 cells transfected with the indicated TrkA constructs (see Figure 3A) and incubated with or without NGF in the presence the cross linker BS3. Molecular weights are indicated at left. Actin was assayed as a loading control.

We considered that, if NGF indeed induces contacts between these eJTM regions, then we should be able to mimic this activity of NGF by forcing the dimerization of eJTMs in the absence of NGF. For this purpose, we individually mutated most of the residues in the eJTM of TrkA to cysteine and subsequently analyzed the dimerization of these transfected single point mutants (Figure 4A). After transfection of Hela cells, disulfide dimers were spontaneously formed in all constructs but the amount of dimers differed between the various mutants (Figure 4B). The amount of dimer is significantly higher in the positions D412C and K411C. As a functional assay we then transfected these mutants into HeLa cells, which do not express endogenous TrkA, and quantified the phosphorylation of the tyrosines from the kinase activation loop (Y674/675) in the absence and presence of NGF (Figure 4C). This analysis showed the presence of active dimers (D406C, K410C and K411C) which are activated constitutively in the absence of NGF, and dimers that are not active in the absence of NGF (V408C, D412C, E413C and T414C). In the presence of NGF these mutants showed no further activation by NGF (Figure 4C) suggesting they are fully active and the dimer interface adopted by the cysteine dimers is similar or identical to the one obtained with NGF binding. In general as a whole there is a poor correlation (red line, R^2^=0.07, Figure 4D) between the amount of dimer formation and constitutive activation, suggesting that dimerization by itself is not enough for TrkA activation. However the mutants with higher constitutive activation (D406C, K410C and K411C) showed a good correlation between dimer formation and activation (Figure 4D, green line, R^2^=0.92). We then transfected some of the active mutants in PC12nnr5 cells. In the absence of NGF the R405C, K410C and K411C mutants induced the formation of neurites in PC12nnr5 cells (Figures 4F and 4G) supporting the constitutive activation of these mutants and suggested that disulfide bond formation through this interface mimics the binding of NGF. If we assume that the TMD α-helix continues in the juxtamembrane region the residues R405, K410 and K411 are in one face of the helix (in green in the Figure 4H). By contrast the residues whose mutation to cysteine do not activate constitutively the receptor are located in another face (in red in the Figure 4H). All the mutants are correctly expressed at the plasma membrane as found by flow cytometry (Figure 4E).

**Figure 4.**
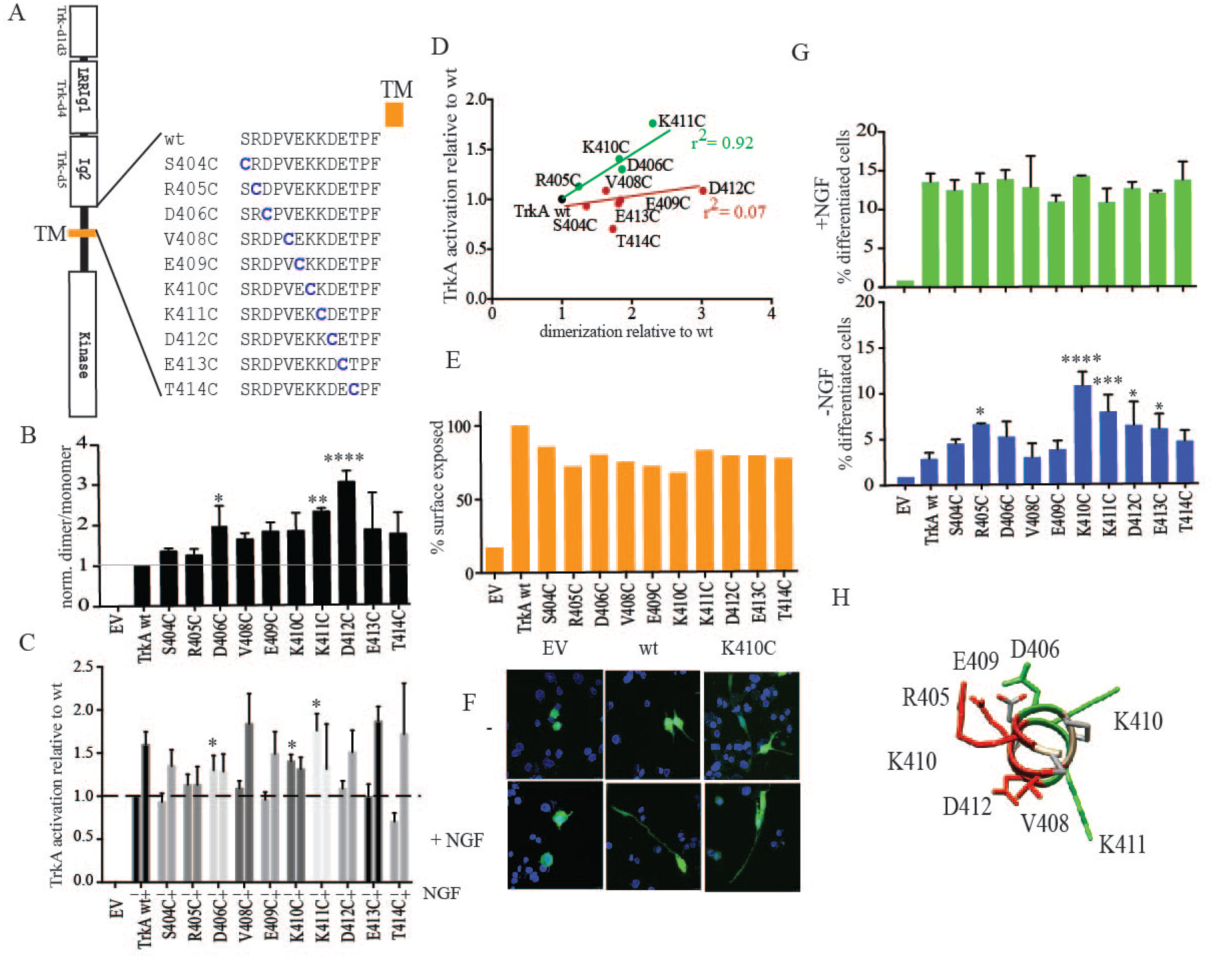
A preferred dimer interface in the TrkA juxtamembrane region. A) Amino acid sequence of the rat TrkA cysteine mutant constructs that are mutated in the region of the eJTM closest to the TMD. B) Quantification of the ratio of dimer:monomer of the TrkA mutants as determined using non-reducing SDS-PAGE. C) Quantification of the activation of TrkA (with and without NGF) by quantifying the signal from the phosphorylation of the Tyr 674/675 signal using western blotting of the cysteine mutants in the JTM region. D) Scatter plot of the dimerization of TrkA cysteine mutants respect to its activation in the absence of NGF. A regression fit using the active (green) and inactive (red) cysteine mutant dimers is shown with the indicated r^2^. E) Cytometric analysis of the expression of the TrkA mutants at the plasma membrane of Hela cells. F) PC12nnr5 cell differentiation assay of TrkA-wt and TrkA-K410C in the presence and absence of NGF. G) Quantification of the differentiation of PC12nnr5 cells transfected with the indicated TrkA constructs and incubated in the absence (blue bars) or presence (green bars) of NGF. H) Model of the eJTM into an ideal α-helix showing the spacial location of the indicted residues. Error bars represent the standard error of the mean. Statistics was done using two-way ANOVA analysis and Dunnet’s multiple comparison test using GraphPad software. The P values of significant differences are shown (**** P<0.0001).

In summary, our results supports the notion of the insufficiency of TrkA dimerization alone for receptor activation and support the existence of an preferred active dimer interface that is formed upon ligand-binding.

### Insertion of leucine residues into the TMD constitutively activates TrkA

It is known that overexpression of TrkA induce a ligand-independent activation of TrkA. The western blot shown in Figure 5A shows that although the overexpression of TrkA induces a ligand-independent activation (probably by facilitating receptor dimerization), the presence of NGF is required for a higher and complete activation of the receptor. This could be the result of an stabilization of the dimer or to a conformational change in the intracellular region of TrkA induced by the binding of the ligand or to both. The complete sequence of the events caused by the ligand is also of great importance for receptor activation. As we have shown above, ligand binding brings eJTM regions into proximity at certain specific positions. In turn, such proximity can either result in a mutual rotation of the receptor TMDs or change the spacing between the N-termini of the TMD helices in the dimer. We found two dimer interfaces in TrkA-TMD that reside on opposite sides of the α-helix suggesting the possibility of a rotation from one interface to the other upon NGF binding. If this model were correct, mutual rotation of two motifs on the surface of the TM helix should activate TrkA in the absence of NGF or increase its basal activation by overexpression. To test this mechanism we introduced a different number of leucine residues into the TrkA-TMD and analyzed the resulting TrkA activation in the absence of NGF (Figure 5B). The insertion of each Leu should rotate the intracellular region around an angle of approximately 100°. Thus, Leu insertion allows evaluation of whether a change in the rotation angle of the intracellular domain plays any role in TrkA activation (Figure 5C). We transfected the constructs TrkA-ins1L, TrkA-ins2L, TrkA-ins3L and TrkA-ins4L that included 1 to 4 inserted leucines respectively. The insertion of one Leu, TrkA-ins1Leu, significantly increased both the constitutive activation of TrkA in transfected Hela cells compared to that of transfected TrkA-wt (Figure 5D and 5E) and the differentiation of PC12nnr5 cells compared to wt-transfected cells (Figure 5F) in a ligand-independent manner. Equal levels of all constructs were expressed at the plasma membrane as determined using flow cytometry (Figure S8). These data indicate that a rotation of the TMD is likely to underlie TrkA activation.

**Figure 5.**
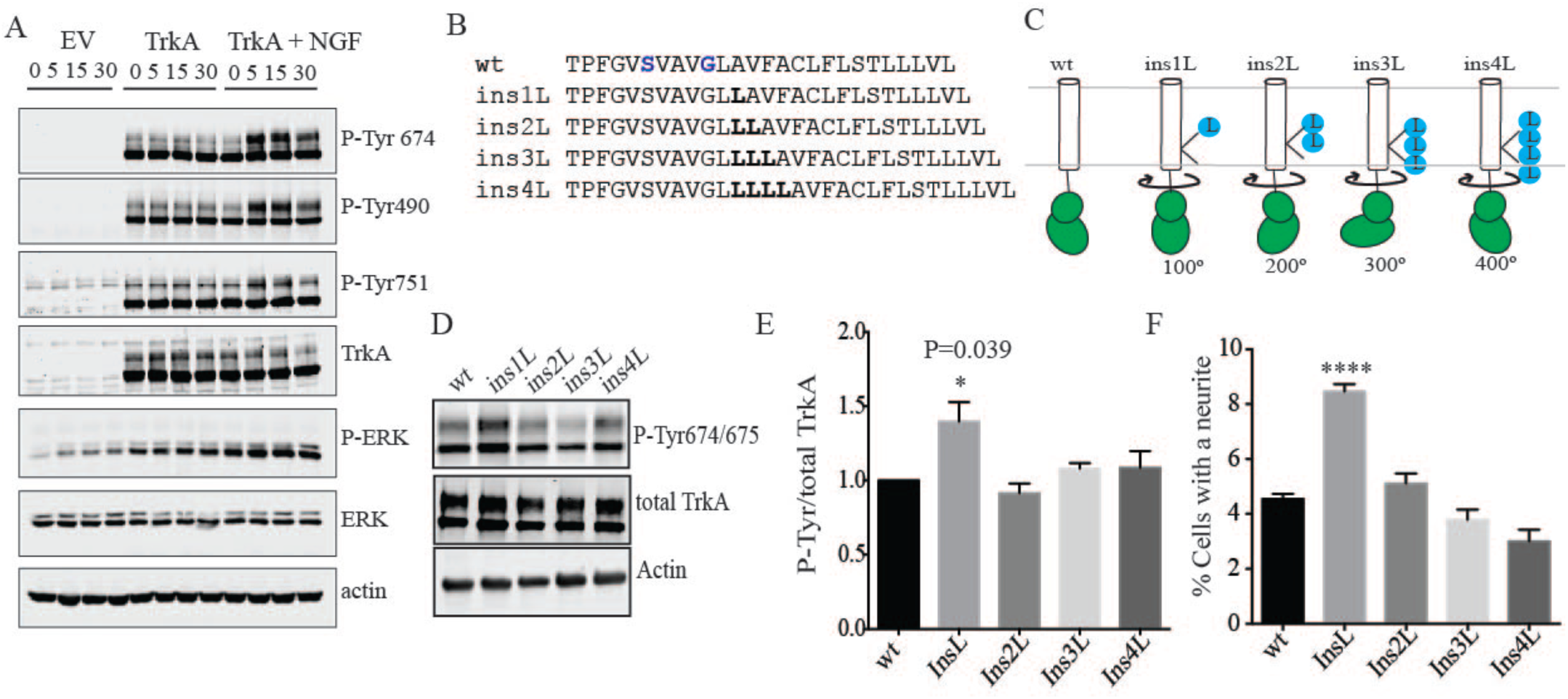
Rotation of the TrkA-TMD as a mechanism of TrkA activation by NGF. A) Effect of NGF on the activation of overexpressed TrkA. Hela cells transfected with the indicated constructs were stimulated with NGF (10 ng/mL) for 0, 5, 15 or 30 minutes. The immunoblots show the activation of TrkA as detected using an antibody specific for the autophosphorylated P-Tyr674/675, P-Tyr490 and P-Tyr751 residues and for the downstream activated P-ERK. The levels of total TrkA and total ERK are shown below each blot. Actin was blotted as a loading control. A representative blot of at least three independent experiments is shown. B) Amino acid sequences of the rat TrkA Leu insertion mutants indicating the location of the inserted Leu residues. C) Schematic drawing of the TrkA-TMD-ICD showing how the different numbers of inserted Leu residues (blue) in the TMD induce a different rotation of the intracellular domain (green). D) Hela cells were transfected with the indicated TrkA-TMD insertion mutants. The immunoblots show the activation of TrkA as detected using an antibody specific for the TrkA autophosphorylation P-Tyr674/675 residues. The levels of total TrkA are shown below. Actin was assayed as a loading control. E) Quantification of the data in D) from at least three independent experiments. Bars represent standard error of the mean. Statistical analysis was performed using one-way ANOVA with Dunnet’s multicomparison test and TrkA-wt as a control. The P value of the significant difference is shown. F) PC12nnr5 cell differentiation of TrkA-TMD insertion mutant-transfected cells incubated in the absence of NGF. The percentage of the total GFP positive-transfected cells with a neurite twice as long as the length of the cell body was quantified. Bars represent the standard error of at least four independent experiments. Statistical analysis was performed using one-way ANOVA with Dunnet’s multicomparison test and TrkA-wt as a control. The P values of significant differences are shown.

## Discussion

TrkA belongs to a subfamily of RTKs that includes the other family members TrkB and TrkC. These RTKs are essential for the formation of the nervous system and mediate a variety of cellular responses in normal biological processes and in pathological states (39). An understanding of their mechanism of action is necessary to facilitate the design of new pharmacological agents targeted to the processes in which they play a role.

Although the general consensus is that Trks are in an equilibrium monomer-dimer, the oligomer state of the Trk receptors in the absence of ligand has been the focus of intense recent research. Using *in vivo* crosslinking and fluorescence complementation assays in PC12 cells that express TrkA, Maruyama et al. found that TrkA, as well as TrkB, form inactive dimers in the absence of NGF (18,40). Marchetti et al., using single-tracking of TrkA receptors, recently found that only around 20% of plasma membrane-expressed TrkA receptors are in a pre-dimer state in the absence of NGF (19). Additionally, the existence of pre-formed dimers of full-length TrkA expressed in the plasma membrane of Xenopus oocytes was shown by freeze-fracture electron microscopy (17). A recent study has provided a further evidence for the existence of pre-formed dimers in the Trk family (20). The laboratory of Dr. Hristova found that TrkA, as well TrkB and TrkC, exist in an equilibrium monomer-dimer state in the absence of ligand depending on the expression levels (20). The authors suggested that the equilibrium constant of TrkA dimerization indicated that endogenous levels, for instance in PC12 cells, the most predominant form of TrkA is a monomer (80%) in equilibrium with a 20% of pre-formed inactive dimers, supporting the data from Marchetti et al. (19). That study also showed that the TMD and the intracellular domains of TrkA contribute significantly to its unliganded dimerization. In summary, although the existence of an equilibrium monomer-dimer(inactive) could be a general assumption for the Trks, in the case of TrkA most of the receptor is in a monomer state.

In this regard, in the present work we posed two major questions: (1) what are the conformations of TrkA TMDs in the active/inactive states? and (2) how is the coupling between the ligand binding and receptor activation? To answer the first question we employed structural characterization using NMR spectroscopy together with mutagenesis studies, disulfide cross-linking and functional assays. We described the high-resolution NMR structure of the TrkA transmembrane domain dimer, which is the first such description of a neurotrophin receptor of the Trk family. The obtained spatial structure was verified using functional assays and crosslinking, which confirmed both the relevance of this structure for TrkA activity and assignation of the found dimer conformation to the receptor active state. This result, combined with the crystal structure of the extracellular domain of the TrkA complex with NGF (16,41), the crystal structure of the TrkA inactive kinase domain (21,22) and the recently reported structures of the entire TrkA intracellular region (42,43) provides an almost complete picture of the full length Trk receptor family, lacking only the structure of the small JTM regions.

Additionally, we found another possible TMD dimerization interface using a genetic assay, and functional tests indicated that this interface was also relevant for TrkA activation. Although destruction of this interface by replacement of a small amino acid residue with a bulky residue inhibited the dimerization in the bacterial membranes, it does not affect the activation of TrkA by NGF. The structural data indicates that this interface is in an opposite side as the active dimer interface. Interestingly in an ideal α-helix connected to the TMD the residues from the eJTM that form inactive dimers when mutated to cysteine are located in the same helix side than the SxxxG motif. Based on this it is possible that the SxxxG interface forms part of the described inactive pre-dimer state, although we cannot detect the formation of dimers when the SxxxG motif is mutated to cysteine in the context of full-length receptor. One reason for this could be that, in the full-length receptor, the distance between the SxxxG motifs are longer than the required for the disulfide bond formation. This could be due to the presence of the soluble domains of TrkA in the full-length receptor that are absent in the bacterial ToxRed constructs. However the finding that the mutation of the G423I reduces significantly the differentiation of PC12 cells with NGF but not the activation of the kinase domain (as shown by the phosphorylation of the Tyr-674/675) is intriguing. PC12 cells differentiation needs of a sustained MAPK activation to differentiate and it is plausible that a higher activation of TrkA is reached formation of oligomers. Actually the formation of TrkA oligomers by NGF has been already described by single-particle tracking in neuroblastoma cells (19). The important role of G423 in cell differentiation and its location in an opposite side of the α-helix to the active dimer interface, suggests that G423 may participate in a protein-protein interface corresponding to a supra-dimer (oligomer) form of the TrkA receptor that is populated upon NGF binding (19). In this context further investigation on the TrkA/NGF oligomer formation is warranted.

To answer the second question posed above, we investigated the role of the eJTM regions of the TrkA receptor upon NGF binding in receptor activation and dimerization. We showed that full-length TrkA receptors can be activated by specific single-point cysteine mutations in the eJTM, in which the position of the mutation relative to the TMD was more important for activation than the dimerization propensity of the mutant. In addition the insertion of Leu residues downstream of the TMD dimerization motif activated TrkA in the absence of ligand. This suggests that rotation of the downstream domain may be behind the activation of the kinase domain. The rotational mechanism of RTK activation has been suggested by other authors (44,45). In this model the ligand would induce a rotation of the TMD dimer interface that will re-orient the kinase domains in order to facilitate the trans-phosphorylation. Although our results support this model of activation other alternative possibilities may exist. For instance the insertion of extra-residues increases the TMD length and could induce a piston-like mechanism of activation by a pulling-down force. However an increase in the length by two, three or four residues should also activate the kinase and this was not the case, as only the Ins1L mutant showed activation. Also as the insertion of the residues are located into the TMD it may alter the dimer interface of the TMD dimer leading to the formation of another dimer interface compatible with a higher activation of the kinase domains. Although we cannot discard this possibility, in the constructs we made the Leu residues are inserted upstream of the active dimer interface in order to not alter the dimer interface found by NMR studies. In any case our data suggest that TrkA requires a reorganization of the JTM and/or TMD regions to couple the ligand binding to kinase activation. This would be of special interest to explain how the binding of the ligand could activate an inactive pre-dimer.

Bearing all of these findings in mind, we propose a mechanism of receptor activation that suggested a ligand-induced dimerization accompanied by some conformational changes in the JTM and TMD regions. This model is supported by our data that showed that cysteine mutants in the eJTM in some specific positions can activate TrkA without ligand, NGF binding brings the N-termini of the TMD α-helices into close proximity and an induced rotation of the TMD generates active molecules of TrkA.

Our model contrasts with the model of TrkB activation recently proposed (46). In that model TrkB is activated by its ligand BDNF and signal in a monomeric state and only after internalization in the endosomes forms dimers (46). Interestingly, and in favor of our model, using the same methodology the authors found that TrkA form dimers with NGF at the plasma membrane. One reason of the different behavior between TrkA and TrkB may reside in the different protein sequences of their TMDs (Figure 1D) that may allow TrkB to reside in a different membrane region or interact with other membrane proteins.

In summary we have provided functional and structural evidences of the roles played by the JTM and the TMD in TrkA dimerization and activation by NGF.

## Material and Methods

### DNA constructs

A plasmid encoding rat TrkA with an N-terminal HA tag was kindly provided by Dr. Y Barde. All TrkA mutants and constructs were derived from this plasmid. Mutagenesis was done using the Site-directed mutagenesis kit (Agilent) according to the manufacturer’s protocol. The oligonucleotide sequences of all of the constructs are available upon request. All DNA constructs were sequenced using local facilities.

### Cell culture and transfection

Hela cells, which do not express endogenous TrkA, were cultured in DMEM medium (Fisher) supplemented with 10% FBS (Fisher) at 37 °C in a humidified atmosphere with 5% CO2. PC12 and PC12nnr5 cells were cultured in DMEM with 10% FBS and 5% horse serum (Fisher). Transfection of Hela cells was performed using polyethylenimine (PEI; Sigma) at a concentration of 1-2 μg/μl. The use of PEI as the transfection reagent for Hela cells resulted in suboptimal transfection (10-15% of cells transfected) and in the expression of only a small amount of TrkA in the cells. In contrast, when the same PEI/DNA ratio was used for transfection of Hek293 cells TrkA was expressed in higher amounts and ligand-independent activation of TrkA was seen. A concentration of 500-1000 ng of DNA per 10 cm cell plate was used for the TrkA activation experiments. Twenty four hours after transfection the cells were <u>lifted</u> and re-plated into 12-well plates at a density of 100,000 cells per well. By using this procedure the percentage of cells transfected was identical in all the wells. Forty eight hours after transfection the cells were starved in serum free medium for 2 h and were then stimulated with NGF (Alomone) at the indicated concentrations and time intervals. Cells were lysed with TNE buffer (Tris-HCl pH 7.5, 150 mM NaCl, 1 mM EDTA) supplemented with 1% triton X-100 (Sigma), protease inhibitors (Roche), 1 mM PMSF (Sigma), 1 mM sodium orthovanadate (Sigma), and 1 mM sodium fluoride (Sigma). In the experiments involving the TrkA cysteine mutants, 10 mM iodoacetamide (Sigma) was added to the lysis buffer. Lysates were kept on ice for 10 minutes and centrifuged at 12,000 g for 15 minutes in a tabletop centrifuge. The protein level of the lysates was quantified using a Bradford kit (Pierce) and lysates were analyzed by SDS-PAGE

### Western blot analysis

Cellular debris was removed by centrifugation at <u>12,000 g</u> for 15 minutes and the protein level of cell lysates was quantified using the Bradford assay (Pierce). Proteins were resolved in SDS-PAGE gels and transferred to nitrocellulose membranes that were incubated overnight at 4 °C with one of the following antibodies: mouse monoclonal anti-HA (1:2000, Sigma); rabbit polyclonal MBP-probe (1:1000, Santa Cruz); rabbit anti-phosphoTyr674/5 (1:1000, Cell Signaling); rabbit anti-TrkA (1:1000, Millipore). Following incubation with the appropriate secondary antibody, membranes were imaged and bands quantified using enhanced chemiluminescence and autoradiography.

### ToxRED assays

Bacterial colonies were inoculated into LB medium (with 50 µg/ml ampicillin). Fresh LB cultures (with 50 µg/ml ampicillin) were inoculated from fresh plates and were grown at 37 °C until they reached an O.D. of approximately 600≈0.6. They were then harvested by centrifugation. Cells were subsequently lysed in lysis buffer (TNE, see recipe above) under gentle conditions to avoid mCherry denaturation. After transfer to 1.5-mL Eppendorf tubes, the mixture was incubated at room temperature with gentle agitation for 30 min. Samples were then centrifuged for 10 min at 12000 × *g* to remove cell debris and to clarify the supernatants for analysis. Aliquots (150 µl) of clarified supernatant were transferred to black, opticlear 96-well plates and mCherry emission spectra were collected using a plate reader (Tecan, Maennedorf, CH) with an excitation wavelength of 587 nm and emission wavelengths of 610–650 nm. Subsequently, aliquots were transferred from black to clear 96-well plates and the absorbance was measured from 450 to 750 nm. Measurements of mCherry and of construct expression were performed from at least 10 different colonies and were normalized for the relative expression level of each construct using western blotting with the anti-MBP antibody. For western blots, samples were mixed with equal volumes of 2× SDS-PAGE sample buffer, heated to 95 °C for 10 min, separated on 10% (w/v) polyacrylamide mini-gels, and were blotted onto nitrocellulose membranes that were probed with anti-MBP antibody. To analyze disulfide bond formation bacterial colonies were lysed using TNE plus 10 mM iodoacetamide and were analyzed using non-reducing SDS-PAGE.

### TrkA-TMD constructs for cell-free expression

The gene encoding the transmembrane domain of the human TrkA-TM (MK_410_KDETPFGVSVAVGLAVFACLFLSTLLLVLNK<u>A</u>GRRNK_447_) was amplified by PCR from six chemically synthesized oligonucleotide templates (Evrogen, Russia) whose sequences partially overlapped along its sequence. The PCR products were cloned into a pGEMEX-1 vector by three-component ligation using the NdeI, AatII and BamHI restriction sites.

### Cell-free gene expression

A bacterial S30 cell-free extract was prepared from a 10 L culture of the E. coli Rosetta(DE3)pLysS strain according to a previously described protocol. The S30 cell extract was stored in 500 μl aliquots at −80 °C. The continuous exchange mode of preparation using a 12.5 kDa membrane was used in this study. Preparative-scale reactions (2-3 ml of reaction mixture) were carried out in 50 mL tubes. Optimal reaction conditions such as Mg2+ and K+ concentrations, the ratio of the reaction mixture (RM) to the feeding mixture (FM) or the DNA concentration were established using homemade reactors based on the Mini-CECF-Reactor previously described (47,48). The final reaction mixture was: a standard FM:RM ratio of 8:1 and a cell-free reaction mixture containing 100 mM HEPES ⁄0.83 mM EDTA with KOH added to achieve a pH of 8.0, 0.1 mg/mL Folinic acid, 20 mM acetyl phosphate, 1.2 mM ATP, 0.8 mM each of G/C/UTP, 2 mM 1,4-dithiothreitol, 0.05% sodium azide, 2% PEG-8000, 20 mM magnesium acetate, 270 mM potassium acetate, 60 mM creatine phosphate, 1 mM each of 20 amino acids or 0.25% of a 20 amino-acid mix (CIL, USA), 1 tablet / 50 mL of complete protease inhibitor (Roche, Switzerland), 0.5 mg/mL E.coli tRNA (Roche, Switzerland), 0.25 mg/mL creatine kinase from rabbit muscle (Roche, Switzerland), 0.05 mg/mL T7 RNA polymerase prepared using a previously described protocol (49), 0.1 U/μL Ribolock (Fermentas), 0.02 μg/μL plasmid DNA, and 30% S30 cell-free extract. All reagents were provided by Sigma unless otherwise specified. Plasmid DNA was purified using a Promega MaxiPrep kit. Reactions were conducted overnight at 34 °C and in an Innova 44R shaker (New Brunswick) set at 150 rpm.

### Protein purification

The cell-free reaction mixture was diluted three-times with buffer A (50 mM Tris pH 8.0 and 200 mM NaCl). After 10 minutes of incubation the mixture was centrifuged for 10 min at 18000 *g* at room temperature. The precipitate was washed consecutively with buffer A containing 30 μg/ml RNAse A (Fermentas) and buffer B (50 mM Tris pH 8.0 and 100 mM NaCl). The target protein was solubilized with 200 μl buffer B containing 1% lauryl sarcosine. After each step the protein was centrifuged for 10 min at <u>18.000</u> *g* at room temperature and aliquots of the supernatant were analyzed using 12,5% Tricine SDS-PAGE (53). The clarified protein solution was applied onto a 10/300 Tricorn column prepacked with Superdex 200 (GE Healthcare) and pre-equilibrated with buffer B containing 0.2% lauryl sarcosine. Protein-containing fractions were combined and precipitated using the TCA/acetone procedure (50).

### Preparation of NMR samples in a membrane mimetic media

The so-called “isotopic-heterodimer” (1:1 mixture of unlabeled and _15_N/^13^С-labeled peptides) samples were prepared corresponding to the TrkA-TMD construct in order to solve its structure. The powder containing the peptides of both samples was first dissolved in a 1:1 trifluoroethanol–H_2_O mixture with the addition of deuterated DPC (d_38_, 98%, CIL) and phosphate buffer, and was then kept for several minutes in an ultrasound bath and lyophilized. Subsequently, the dried samples were dissolved in 350 µl of a 9:1 H_2_O:D_2_O mixture. To attain a uniform micelle size and uniform distribution of the peptide throughout the micelles, the samples were sonicated in an ultrasound bath for several minutes until the solution was completely transparent. The TrkA-TMD concentration in the isotopic-heterodimer sample was 1.9 mM, and other conditions were: LPR 50:1, pH 5.9, and 20 mM phosphate buffer. Samples were placed in Shigemi NMR tubes with a glass plunger. Selective-residue labeling was implemented to avoid peaks overlapping while processing the NMR spectra.

### NMR spectroscopy and spatial structure calculation

NMR spectra were acquired at 45 °C using 600 and 800 MHz AVANCE III spectrometers (Bruker BioSpin, Germany) equipped with pulsed-field gradient triple-resonance cryoprobes. 1H, 13C, and 15N resonances of TrkA-TMD were assigned with CARA software (51) using two- and three-dimensional heteronuclear experiments (52): ^1^H/^15^N-HSQC, ^1^H/^15^N-TROSY, ^1^H/^13^C-HSQC, ^1^H/^15^N-HNHA, ^1^H/^13^C/^15^N-HNCA, ^1^H/^13^C/^15^N-HN(CO)CA, ^1^H/^13^C/^15^N-HNCO, 3D-HccH-TOCSY, ^13^C- and ^15^N-edited NOESY-HSQC (recorded on 600 and 800 MHz spectrometers, respectively). Dimeric spatial structures were calculated with the CYANA program (53) based on torsion angle restraints estimated from the chemical shift values obtained with the standard protocol of the TALOS-N program (54) and with intra- and inter-monomeric NOE distance restraints derived through analysis of the three-dimensional ^15^N- and ^13^C-edited NOESY and ^15^N,^13^C-F1-filtered/F3-edited-NOESY spectra (52) acquired for “isotopic-heterodimer” samples. MOLMOL software was used to calculate the contact areas between the dimer subunits and to visualize the structures (55). Hydrophobic properties of the α-helices in the TrkA-TMD dimers were calculated using the molecular hydrophobicity potential (MHP) approach implemented in the PREDDIMER program (56).

### Isolation of Membrane fractions

TrkA mutants were overexpressed in Hela cells by transfection using PEI (1mg/ml). 48 hours after transfection cells were resuspended in 1ml ice-cold homogenization buffer (250 mM sucrose, 1mM EDTA, 10 mM tris buffer pH 7,1 plus protease inhibitors) and broken by sonication in two time intervals (30,30 s) with 50W and frequency at 30 MHz on ice. The broken cell homogenate was centrifugated for 10 minutes at 500g at 4°C to remove whole cells and nuclei. To collect the membrane fraction the cleared supernatant was centrifuged at 100,000*g* at 4°C for 1h in a Beckman Optima MAX ultracentrifuge with a TLA110 rotor using polycarbonate thick wall centrifuge tubes (Beckman coulter ref. 362305). The supernatant that contains soluble proteins was removed and the pellet-containing membranes was resuspended in 1ml ice-cold homogenization buffer by sonication and re-centrifuged at 100,000*g*. The final pellet contains the membrane fraction used to the iodine oxidation protocol.

### Iodine oxidation for crosslinking of TrkA mutants

As cysteine residues are buried deep in the phospholipid bilayer, membranes were isolated and I2 was used as the oxidation agent following as in (Schwem & Fillingame, 2006). Membrane fractions were diluted in the homogenization buffer (250 mM sucrose, 1mM EDTA, 10 mM Tris buffer pH 7,1 plus protease inhibitors). Protein content was quantified using a Bardford kit (Pierce) and equal amounts of isolated membranes fractions were incubated for 10 min with or without NGF (10 ng/mL). A solution of 2.5 mM I2 in absolute ethanol was freshly prepared immediately before the crosslinking of cysteine residues and was added to the incubated membrane fractions with NGF (250 mM final Iodine concentration) for 30 seconds at room temperature. The reaction was stop adding 1/10 volume of a freshly made solution of sodium thiosulfate (60 mM final concentration). Non-reducing SDS-PAGE sample buffer was added and the samples were boiled for 5 minutes before analyzed by non-reducing SDS-PAGE.

### Differentiation of PC12nnr5 cells

Transfection in PC12nnr5 cells was performed using Lipofectamine 2000 as per manufacturer instructions. The mutant or the wild-type TrkA-transfected cells and mock-transfected cells as well as non-transfected cells were treated under the same conditions in a six-well tissue culture plate. The cells were washed three times with serum-free medium and incubated for 48 h in a medium containing 1% fetal bovine serum and 50 ng/mL of NGF (Alomone). At time 0, 24h and 48h, cells were washed with cold PBS and fixed with 4% paraformaldehyde during 15 minutes at room temperature. Cells were imaged using a Leica SP8 spectral confocal microscope. The percentage of cells with a neurite twice longer as the cell body was counted as differentiated.

### Accession codes

Atomic coordinates and experimental restraints were deposited in the Protein Data Bank under the accession codes PDB ID: 2n90 for TrkA-TM-wt.

## Acknowledgments

We thank Dr. Carlos Ibañez for critical reading of the manuscript, Dr. W. DeGrado for providing ToxRed plasmids, and Dr. Yves Barde for the TrkA plasmid. This study was supported by the Spanish Ministry of Economy and Competitiveness (MINECO; project BFU2013-42746-P and SAF2017-84096-R) and by the Generalitat Valenciana Prometeo Grant 2018/055 to MV). Studies of TrkA-TM homodimerization were supported by the Russian Science Foundation (grant No# 19-74-30014 to A.S.A).

## Author Contributions

M.L.F. constructed the TrkA mutants and performed the TrkA activation, differentiation, functional and crosslinking experiments. K.D.N. designed and performed NMR experiments and analyzed NMR data. S.A.G. developed the expression protocols and synthesized samples for NMR analysis. K.S.M. analyzed NMR data and contributed to the preparation of the manuscript. A.S.A. and M.V. designed and supervised the entire project and wrote the manuscript.

---

**Figure 1- Table supplement 1.**
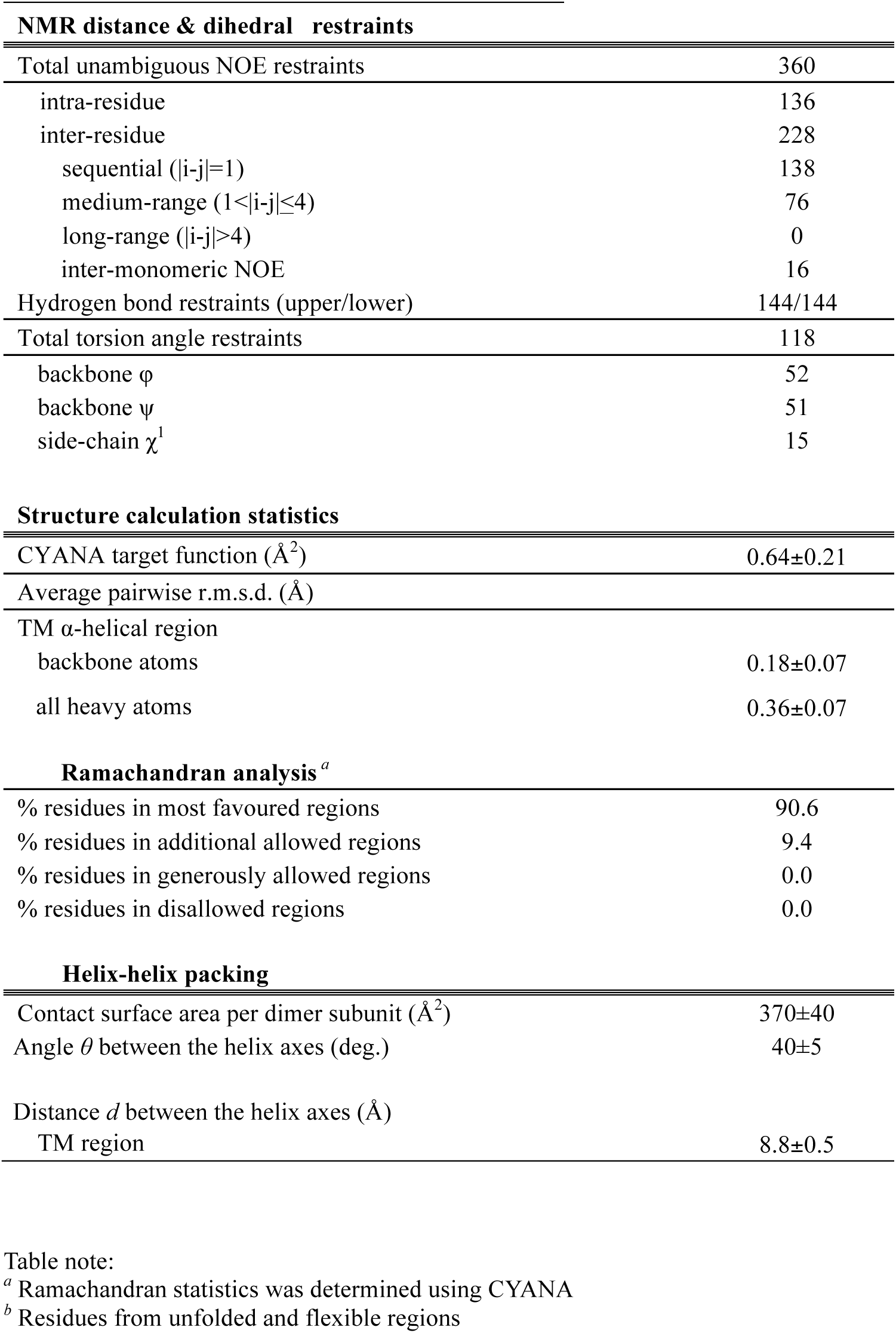

## Supplemental Figure Legends

**Figure S1.**
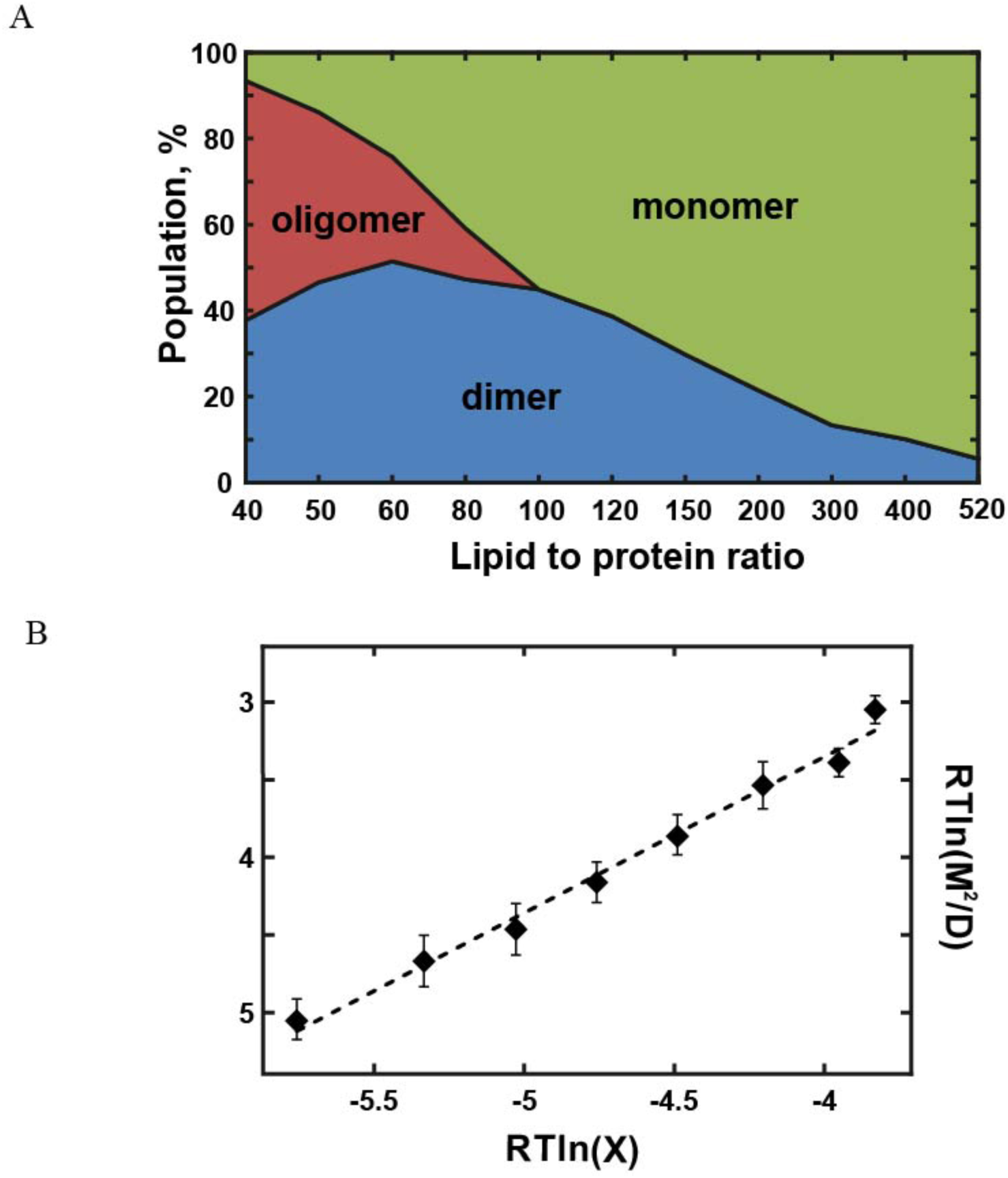
Dimerization of the TrkA-TMD in micelles. (A) Percentage of the monomeric (green), dimeric (blue) and oligomeric (red) forms of the human TrkA-TMD at different LPRs. Population values were calculated from the cross-peak intensities in the ^1^H/^15^N-TROSY spectra. (B) Apparent free energy of dimerization of the TrkA-TMD (ΔGapp)=RTln([M]2/[D]) plotted as a function of RTln(X), where X = ([Det]−Nm[M]−Nd[D])/Ne. [M], [D] and [Det] represent monomer, dimer and DPC concentration, respectively. Ne represents the number of DPC molecules per empty micelle, and Nm and Nd represent the number of DPC molecules per micelle for monomers and dimers, respectively.

**Figure S2.**
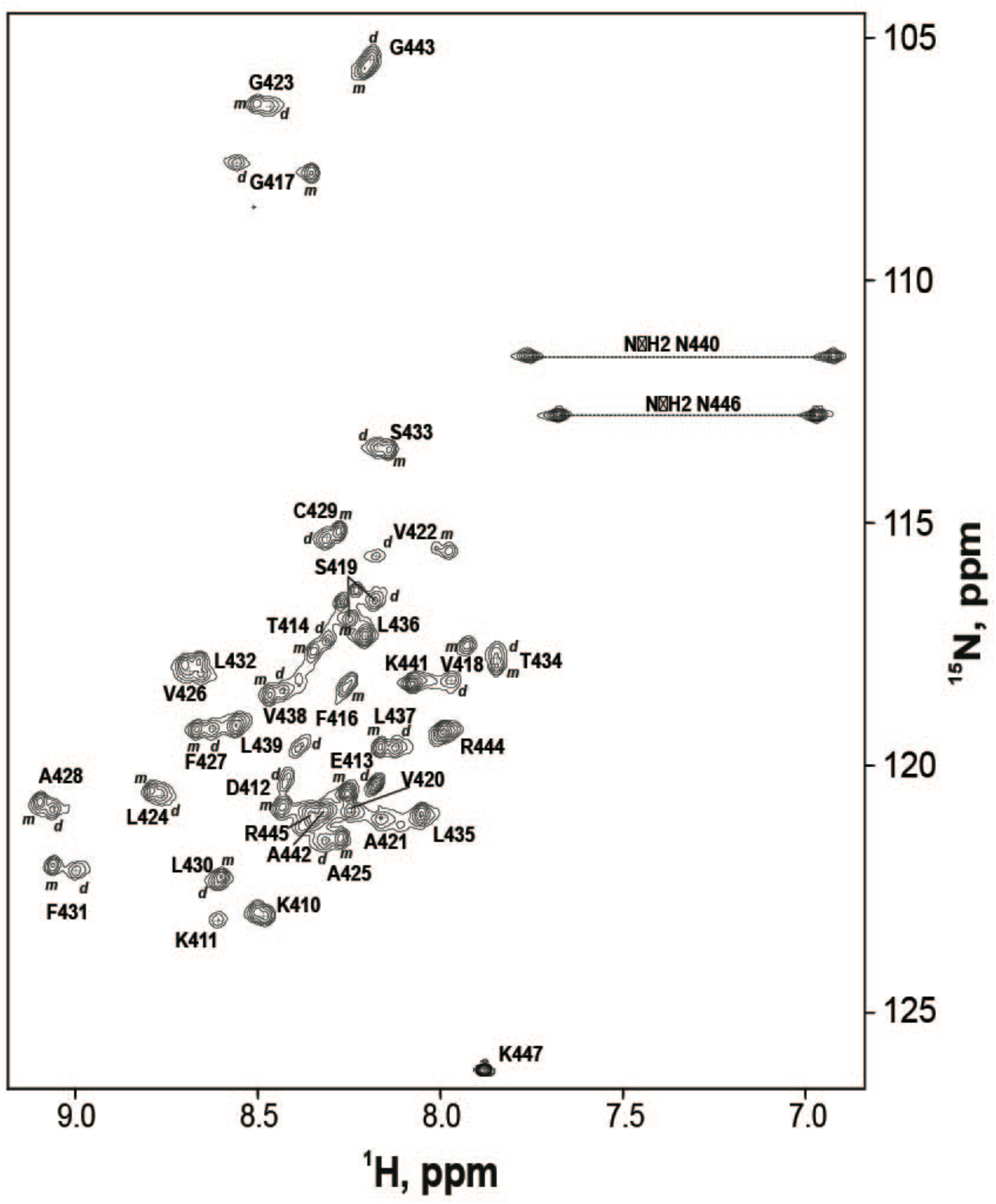
The ^1^H/^15^N-HSQC spectrum of the human TrkA-TMD-wt. The human TrkA-TMD was solubilized in an aqueous suspension of DPC micelles at an LPR of 50:1, and at 45°C and pH 5.9. ^1^H–^15^N backbone and side-chain resonance assignments are shown.

**Figure S3.**
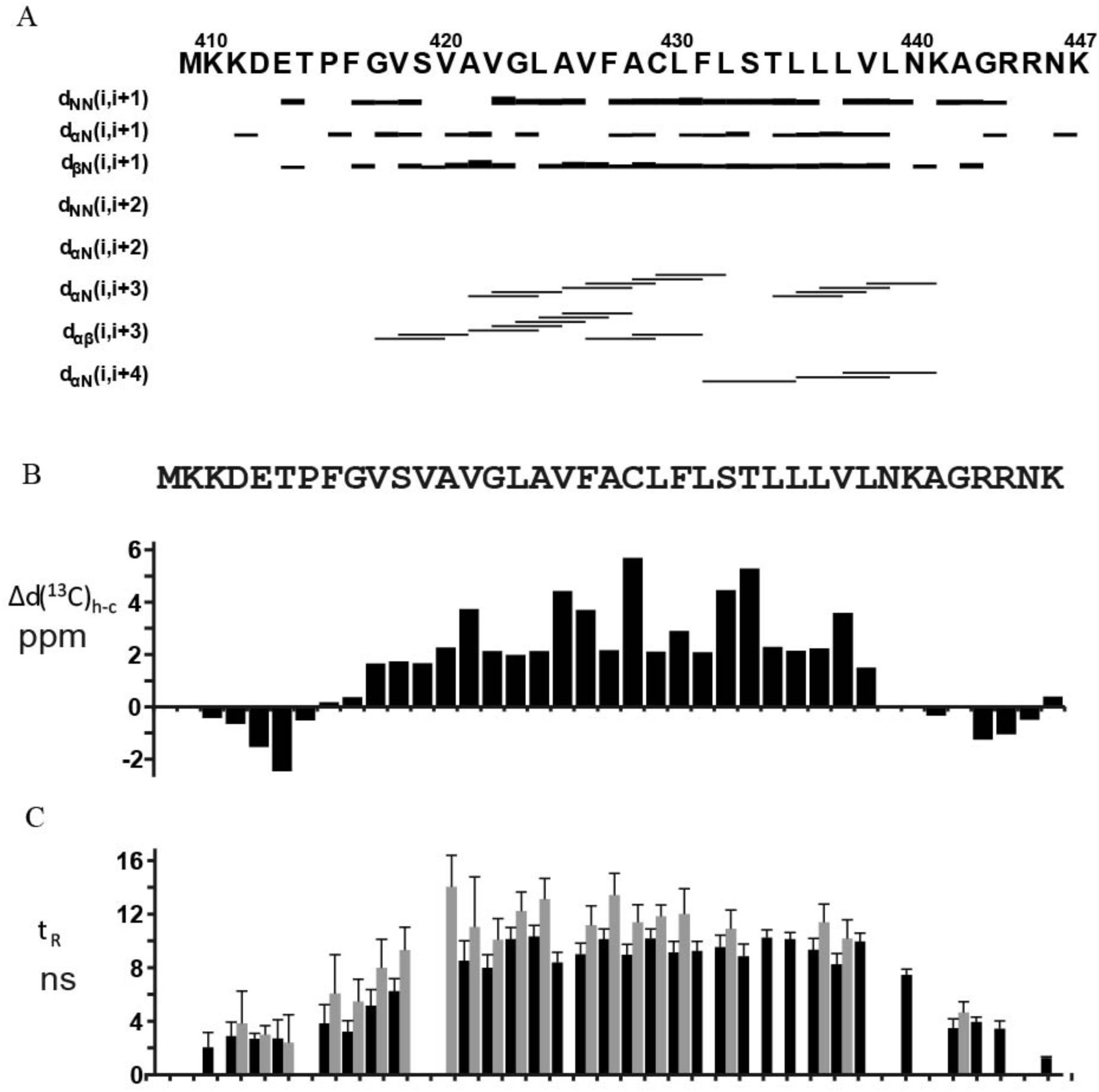
Summary of the NMR data for the TrkA-TMD embedded into DPC micelles. A) NOE connectivities observed in the ^1^H/^15^N-NOESY-HSQC spectrum for the TrkA-TMD peptide. B) Secondary ^13^Cα chemical shifts of the TrkA-TMD are given by the difference between the actual chemical shift and a typical random-coil chemical shift for a specific residue. Pronounced positive or negative Δδ(^13^Cα)h−c values indicate a helical structure or an extended conformation of a protein. C) Local rotation correlation times t_R_ (ns) values of the amide groups derived from ^15^N-relaxation data. Decreased t_R_ values indicate enhanced local flexibility of the protein backbone.

**Figure S4.**
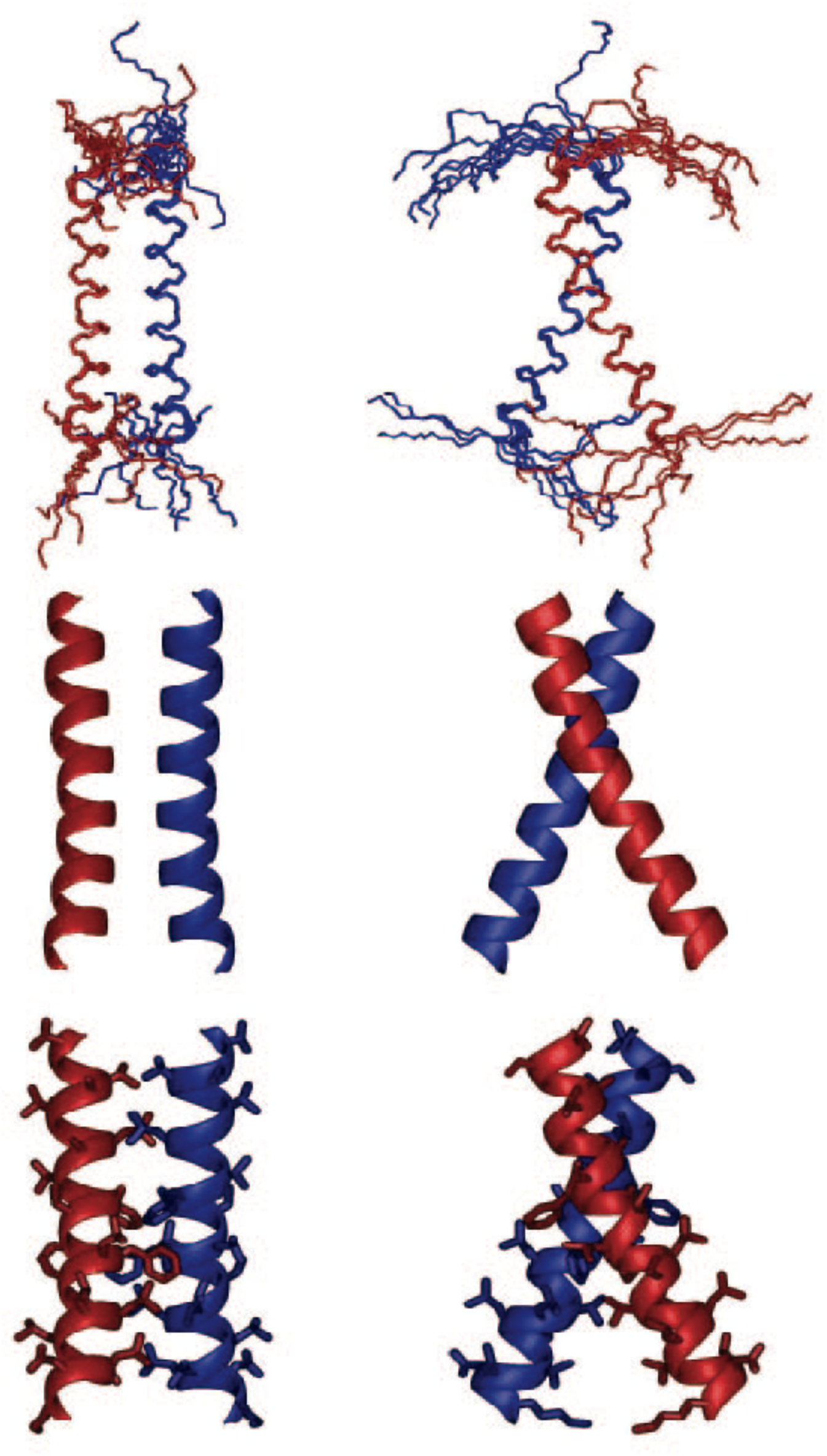
Assessment of the quality of the NMR-derived TrkA-TMD dimer structure. Structural alignment of the 10 lowest energy NMR structures of TrkA-TMD dimers.

**Figure S5.**
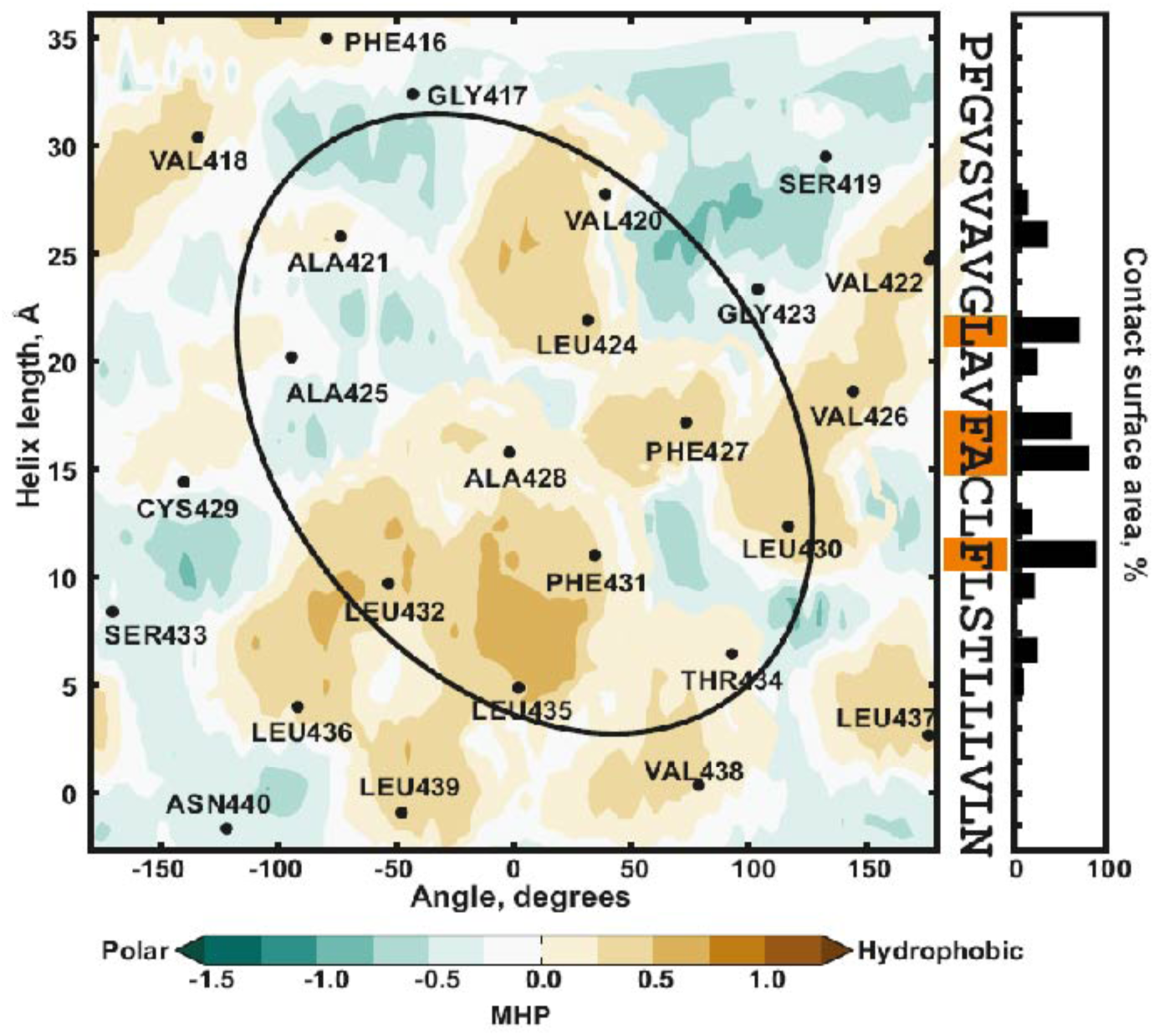
Hydrophobicity map of the TrkA-TMD dimer. The black oval delineates an helix-packing interface as calculated with the PREDIMER software. The residues within this black oval comprise the TrkA-TMD dimerization interface. The panels to the right show the percentage of inter-monomer contact surface area.

**Figure S6.**
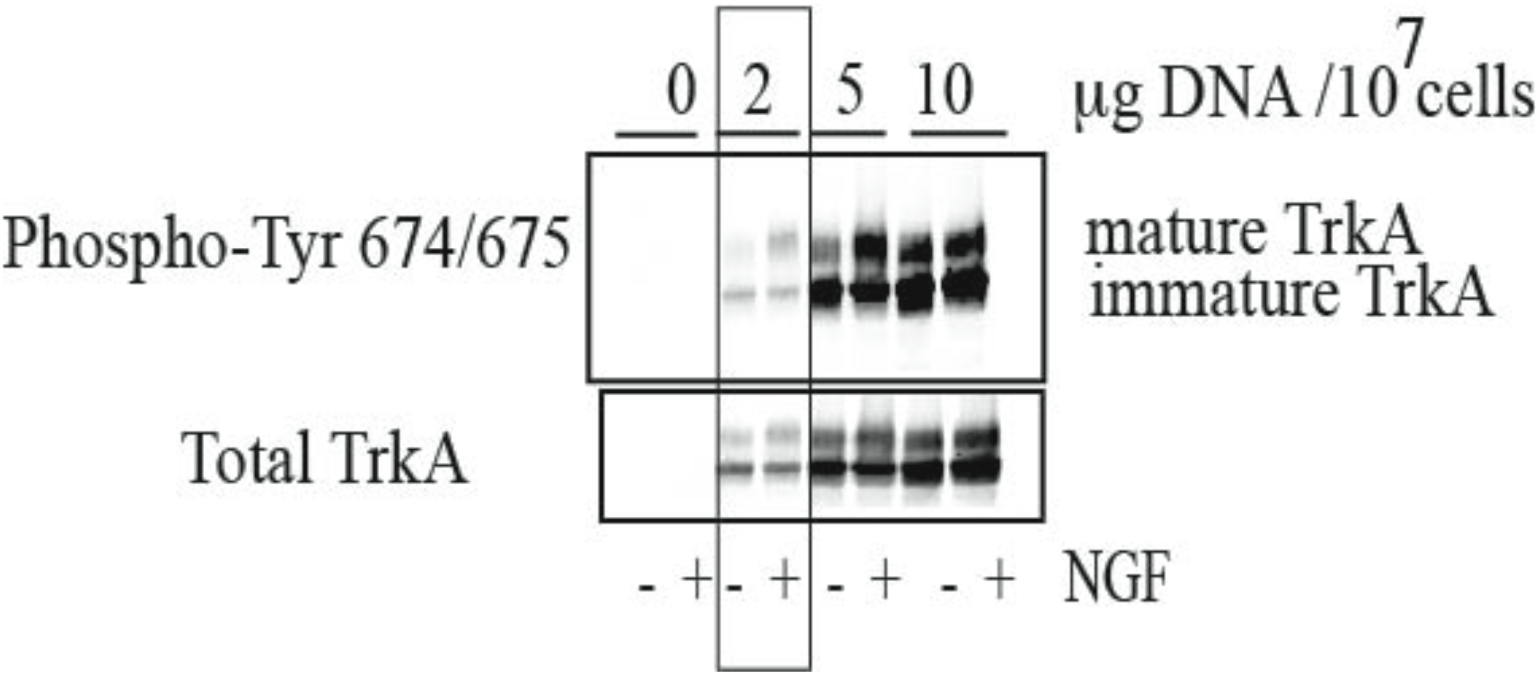
TrkA-concentration-dependence of ligand-independent activation of TrkA. Increasing amounts of TrkA were transfected into Hela cells using PEI. After 48 h, the cells were stimulated or not with NGF (10 ng/mL). Cell lysates were analyzed by SDS-PAGE immnoblotting using antibodies specific for TrkA phospho-Tyr674/675 and total TrkA.

**Figure S7.**
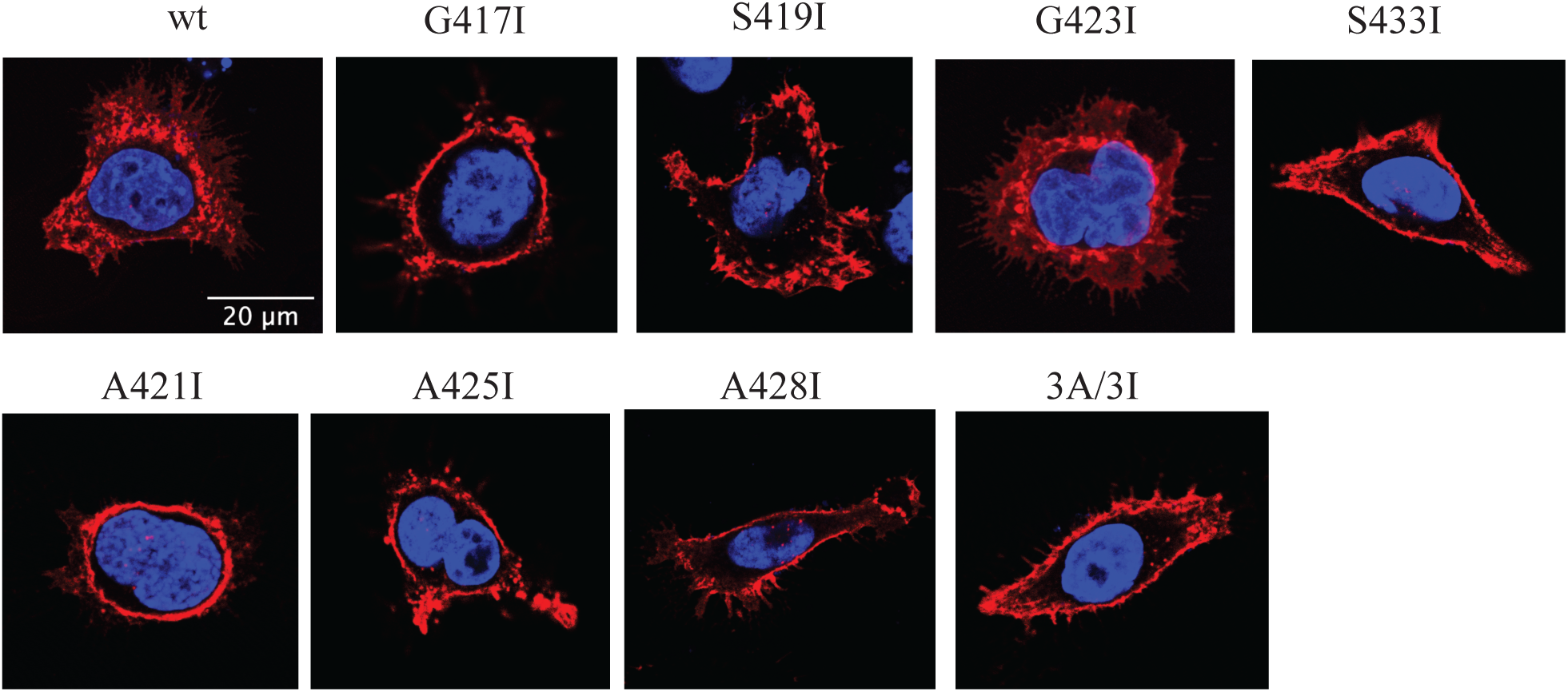
Membrane localization of TrkA mutants in Hela cells. Confocal fluorescence images of Hela cells transiently transfected with the indicated TrkA constructs. The cells were fixed and analyzed using an antibody against HA located in the N-terminus of TrkA, which was detected with a secondary antibody conjugated with the fluorophore alexa-555 (Red). Blue, nuclear DAPI staining. The cells shown are representative of 10 transfected cells analyzed for each condition.

**Figure S8.**
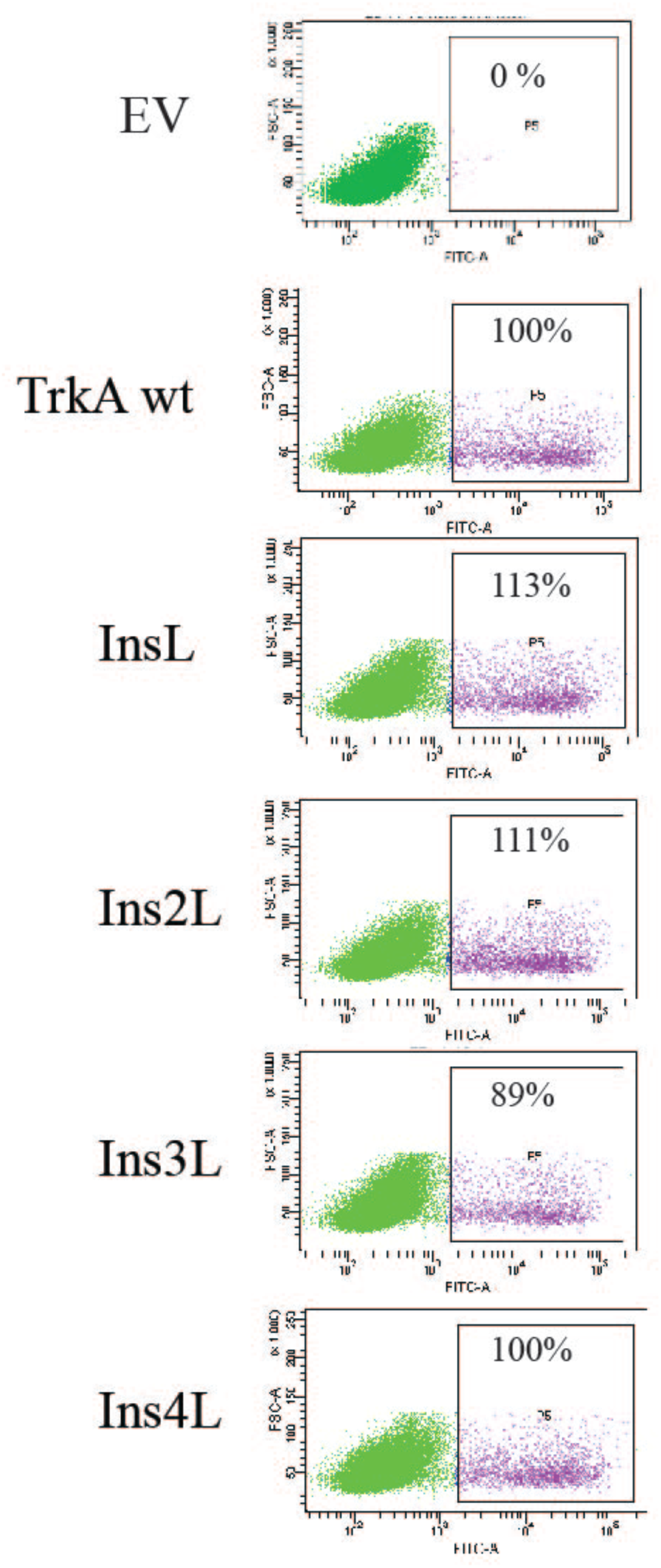
Membrane localization of TrkA mutants in Hela cells by cytometric analysis. Percentage of TrkA constructs expressed at the plasma membrane relative to TrkA-wt (100%) obtained in Hela cells transfected with the indicated constructs and analyzed by flow cytometry using an HA primary antibody and a secondary antibody labeled with a fluorophore for alexa-488. EV is empty vector.

